# Soluble ZP2 N-terminal fragments activate CatSper-dependent Ca^2+^ entry and regulate motility and acrosomal exocytosis in mammalian sperm

**DOI:** 10.64898/2026.08.19.745727

**Authors:** Caroline Wiesehöfer, Lorena Sunjic, Lara Suetin, Jong-Nam Oh, Marc Wiesehöfer, Mykola Lyndin, Thiviya Thayaparan, Layani Chandrakumar, Jean-Ju Chung, Gunther Wennemuth

## Abstract

Sperm motility and function are central to mammalian fertilization and are tightly regulated by intracellular calcium (Ca²⁺) signaling. This signaling is primarily orchestrated by the CatSper Ca²⁺ channel complex located in the flagella of spermatozoa. However, the natural ligands that activate CatSper remain largely unknown in many species, despite the conservation of CatSper in mammals. Here, we present a signaling role for soluble N-terminal ZP2 fragments in regulating CatSper activity and sperm physiology in mice and humans. ZP2 has been implicated in mediating sperm binding and recognition at the oocyte surface interface; however, new evidence is starting to unveil the molecular mechanisms and function of ZP2 during fertilization transition. Here, we show that the during fertilization, cleaved ZP2 N-terminal fragment triggers a rapid and robust increase in intracellular Ca²⁺ levels in sperm. This increase depends strictly on CatSper function, as demonstrated through pharmacological analysis and *CatSper1* knockout mice. This calcium influx is sufficient to induce acrosomal exocytosis in a subset of human and mouse sperm. AlphaFold-based structural modeling suggests a potentially conserved extracellular interaction site between the soluble N-terminal ZP2 fragments and the CatSper complex. In human sperm, ZP2 treatment significantly modulates motility parameters, including flagellar movement and velocity, while inducing a CatSper-dependent increase in intracellular Ca²⁺ similar in magnitude to that evoked by progesterone. Species-matched ZP2 stimulation elicits the stronger calcium response, underscoring evolutionary adaptations in ligand-channel protein pairs. Taken together, our findings reveal a conserved signaling pathway from ZP2 to CatSper that integrates oocyte-derived signals into the regulation of sperm motility and acrosomal exocytosis. This pathway provides new mechanistic insights into fertilization and highlights potential targets.

## Introduction

In mammalian fertilization, the species-specific binding of a spermatozoon to the zona pellucida (ZP), the glycoprotein-rich extracellular matrix of the oocyte, represents a critical checkpoint that must occur before the sperm can penetrate the ZP and fuse with the oocyte. The ZP is composed of four glycoproteins in humans (ZP1-4) (Spargo & Hope, 2003) and three in mice (ZP1-3) (Bleil & Wassarman, 1980), all of which play roles in sperm binding and selection (Ata *et al*, 2026) (Ata *et al*., 2026; Avella *et al*, 2014b; Bleil & Wassarman, 1980; Gupta, 2021; Tian *et al*, 1997). Among these, ZP2 has classically been viewed as a key determinant of sustained binding of acrosome-reacted sperm and species-specific recognition, complementing the initial attachment mediated by other zona components (Chiu *et al*, 2008). Its N-terminal region contains the best-defined sperm-recognition site (e.g. residues 51-149 in human ZP2) and is also crucial for preventing polyspermy (Avella *et al*, 2013). During fertilization, cortical granule exocytosis releases the metalloproteinase ovastacin (ASTL), which cleaves ZP2 at a conserved site in the N-terminus (Burkart *et al*, 2012; Gahlay *et al*, 2010). This cleavage has long been interpreted as functionally switching the zona from “receptive” to “non-receptive”, supported by genetic models in which loss of ovastacin or expression of non-cleavable ZP2 results in persistent sperm binding and polyspermy (Avella *et al*., 2014b; Korschgen *et al*, 2017; Nishio *et al*, 2024b; Xiong *et al*, 2017b). Recent high resolution structural and genetic work (Nishio *et al*, 2024a) refines this model by showing that deletion of the ZP2 N1 domain does not abolish sperm attachment, but instead promotes a semi-hardened zona and subfertility (Nishio *et al*., 2024a). Mechanistically, ZP2 cleavage drives supramolecular oligomerization and crosslinking of ZP filaments to the cleaved N-terminal fragments, thereby rigidifying the matrix as a predominantly mechanical barrier of penetration rather than simply removing a binding site. In this updated framework, the N1 domain in the intact ZP2 can act as a spacer that prevents premature filament packing and may help present the cleavage site to ovastacin (Nishio *et al*., 2024a). Interestingly, a partial cleavage of ZP2 even before fertilization was reported, preventing sperm-ZP binding and fertilization (Nishio *et al*., 2024a). However, the precise function of the soluble N-terminal ZP2 fragments once it is cleaved and released remains unknown.

For a spermatozoon to reach and penetrate the ZP, it must first gain fertilizing potential through capacitation (Austin, 1952; Chang, 1951), a Ca^2+^-dependent process that triggers a powerful, whip-like swimming pattern known as hyperactivated motility (Puga Molina *et al*, 2018; Suarez *et al*, 1993). This critical calcium influx is mediated by the CatSper channel, a complex, sperm-specific ion channel composed of four pore-forming subunits (CatSper1-4) and more than ten auxiliary subunits including the Ca^2+^ sensor, EFCAB9 (Hwang & Chung, 2023; Hwang *et al*, 2019; Vyklicka & Lishko, 2020). Consequently, the disruption of CatSper function, whether through genetic mutation or pharmacological inhibition, dysregulates sperm hyperactivation (Chung *et al*, 2017; Wiesehofer *et al*, 2022; Young *et al*, 2024) and male fertility (Chung *et al*, 2011; Huang *et al*, 2023; Hwang & Chung, 2023; Hwang *et al*, 2022; Qi *et al*, 2007; Quill *et al*, 2003; Ren *et al*, 2001). Beyond driving hyperactivation, CatSper and its auxiliary subunit EFCAB9 are also critical for maintaining the sperm’s typical clockwise, chiral swimming pattern (Wiesehofer *et al*., 2022). Intriguingly, both the loss of CatSper or EFCAB9 function and treatment with the N-terminal ZP2 fragments disrupt this directional movement (Wiesehofer *et al*., 2022). This finding illustrates a potentially functional crosstalk between the oocyte’s matrix-derived components and the sperm’s calcium signaling machinery (Wiesehofer *et al*., 2022).

Understanding the interplay between ZP/ZP-derived components and sperm Ca^2+^ signaling is critical, as this finely tuned system holds the key to both fundamental reproductive biology and the development of novel non-hormonal contraceptive strategies. Reflecting its importance, mutations in the *ZP2* gene can cause abnormal ZP matrix formation and lead to female infertility (Dai *et al*, 2019). This central role has also made ZP2 a promising target for non-hormonal contraception, with research exploring the use of ZP2-derived peptides and specific antibodies to block fertilization (Avella *et al*, 2016; Dioguardi *et al*, 2025). However, a central question remains unanswered: how does the oocyte’s ZP2-N-terminus modulate the sperm’s CatSper channel? While it is known that the entire solubilized ZP can trigger CatSper-mediated Ca^2+^ influx, this effect is indirect and dependent on other signaling events, such as changes in intracellular pH and cAMP levels (Balbach *et al*, 2020a). Therefore, a direct, mechanistic link between a specific ZP protein and CatSper activation has yet to be established, representing a significant gap in our understanding of fertilization.

In this study, we identify a N-terminal ZP2 fragment (not the intact polymeric ZP2 within the zona matrix) as a physiological activator of the sperm CatSper channel. Using both mouse and human models, we show that recombinant soluble N-terminal ZP2 fragments induce a rapid, robust, and CatSper-dependent Ca^2+^ influx in capacitated sperm. This ZP2-driven signal triggers downstream events including sperm motility changes and acrosomal exocytosis, both essential for fertilization. Our structural prediction/modeling combined with motility analyses suggest a potentially evolutionarily conserved signaling axis where soluble ZP2 N-terminal fragments can function as zona-derived cues for the sperm’s CatSper-driven functional program. These results define a fundamental mechanism of gamete communication and sperm activation at the point of fertilization.

## Materials and Methods

### Reagents

Lectin from *Pisum sativum*, A23187, Mibefradil dihydrochloride hydrate, chlortetracycline-hydrochloride and Progesterone (Sigma-Aldrich, Darmstadt, Germany); Vectashield^®^ Plus Antifade mounting medium with DAPI (H-2000, Vector Laboratories, Susteren, Netherlands); Pluronic™ F-127, Fluo-4 AM (Thermo Fisher Scientific, Dreieich, Germany); NNC 55-0396 (Cayman Chemical Company); HC-056456 (MedChemExpress, South Brunswick, New Jersey, USA); Zeocin^TM^ (Invivogen, Toulouse, France).

### Animals

Wild-type C57BL6/J mice were acquired from Charles River Laboratories (Erkrath, Germany). B6D2-Tg(CAG/Su9-DsRed2,Acr3-EGFP)RBGS002Osb (referred to SU9/Acr3) transgenic mice were provided by the RIKEN BioResource Center (Tsukuba, Ibaraki, Japan), and *CatSper1*-null mice (*CatSper1^-/-^*) were obtained from The Jackson Laboratory (Bar Harbor, ME, USA). All animal experiments were conducted in accordance with the institutional guidelines and ethical regulations of the University of Duisburg-Essen and were approved by the responsible state authority (Landesamt für Natur, Umwelt und Verbraucherschutz Nordrhein-Westfalen, LANUV). Animals were housed and maintained in the Central Animal Laboratory (ZTL) of the University Hospital Essen.

### Human sperm donors and human tissue

All experiments involving human spermatozoa were approved by the Ethics Committee of the University of Duisburg-Essen (approval number 14-5748-BO). Written informed consent was obtained forms all donors prior to sample collection, and all samples were pseudonymized to protect donor identity. Donors were instructed to maintain sexual abstinence for 48–72 hours before sample donation.

All ovarian tissue samples in this study were collected and preserved at the Scientific Center of Pathomorphological Research, Sumy State University (Sumy, Ukraine). The use of human resected tissue specimens was approved by the Ethics Committee of Sumy State University (Protocol No. 14/65; Approval Date: March 14, 2025). Written informed consent was obtained from all patients prior to tissue collection. All tissue samples and associated clinical data were fully anonymized prior to analysis.

### Cell lines

HEK293T cells (ATCC/LGC Standards GmbH, Wesel, Germany) were cultured in DMEM (Thermo Fisher Scientific, Oberhausen, Germany) supplemented with 10% heat inactivated fetal calf serum (FCS; Sigma Aldrich, Hamburg, Germany), 100 U/ml penicillin and 100 µg/ml streptomycin (100 µg/ml) (Life Technologies/Gibco, Oberhausen, Germany). These cells were used for recombinant production of murine and human ZP2 proteins. The hybridoma cell lines CRL-2463™ and CRL-2568™ (ATCC®) were used to produce monoclonal antibodies specific to murine ZP2 (aa 114–129) and human ZP2, respectively. Antibody production and purification were performed as previously described (Wiesehofer *et al*., 2022).

### Sperm preparation

Murine spermatozoa were prepared as previously described (Wiesehofer *et al*., 2022). Briefly, cauda epididymis and vas deferens were surgically excised from mice and then rinsed with HS medium (containing in mM: 135 NaCl, 5 KCl, 1 CaCl_2_, 2 MgCl_2_, 230 mM HEPES, 510 mM glucose, 10 DL-lactic acid and 10 pyruvic acid; pH adjusted to 7.4 with NaOH; 353 mOsm). Spermatozoa were allowed to swim out into HS supplemented with 15 mM NaHCO_3_ (HSB) through small incisions for 15 min at 37°C under 5% CO_2_. Capacitation was induced by incubating spermatozoa in HSB containing 5 mg/ml BSA (pH 7.4, 373 mOsm) for 2 h at 37°C and 5% CO_2_. Spermatozoa were washed three times by centrifugation (400 x g, 5 min), and all subsequent procedures were carried out at room temperature (22-25°C) in HS or capacitating medium.

Human spermatozoa were prepared as previously described (von Ostau *et al*, 2024). In short, 1 ml of ejaculate was layered under 4 ml of HS buffer and incubated for 60 min at 37°C and 5% CO_2_, allowing motile spermatozoa to swim up into the upper layer (swim-up method). Motile sperm were collected from the supernatant for subsequent experiments.

### Plasmids

Both, cDNA encoding murine ZP2 (amino acids 35-149) or human ZP2 (amino acids 39-154 aa) was amplified by PCR and cloned into pSEC-Tag2 expression vector (Invivogen, Toulouse, France) to generate muZP2-pSEC and huZP2-pSEC constructs. Primer sequences used for cloning are shown in Table S1.

### Stable Transfection

To produce N-terminal ZP2 peptides, HEK293T cells were transfected with expression plasmid DNA (muZP2-pSEC; huZP2-pSEC or empty pSEC) using Polyfect transfection reagent (Qiagen, Hilden, Germanyaccording to the manufacturer’s instructions. Stable transfectants were selected with Zeocin^TM^ (200 µg/ml; Invivogen, Toulouse, France). Expression of muZP2^35-149^ and huZP2^39-154^ was confirmed by Western blot analysis. Finally, cells were adapted to BalanCD HEK293 medium (FUJIFILM Europe B.V., Tilburg, Netherlands) supplemented with BalanCD HEK293 Feed (FUJIFILM Europe B.V., Tilburg, Netherlands) to replace FCS and increase protein yields.

### Protein purification

Recombinant ZP2 proteins were purified as previously described (Wiesehofer *et al*., 2022). Briefly, cell culture supernatants were sterile filtered, supplemented with 5X binding buffer (in mM: 100 Na_3_PO_4_, 2500 NaCl, 150 Imidazole, pH 7.5) and incubated for 3 h with Ni-NTA-agarose beads (Quiagen, Hilden, Germany), pre-washed with PBS. Beads were washed twice with 1X binding buffer. Bound proteins were eluted by incubation for 30 min in elution buffer (in mM: 20 Na_3_PO_4_, 500 NaCl, 1000 Imidazole, pH 7.2) and beads were removed by centrifugation. The eluate was dialyzed against PBS (pH 7.4) (pore size 4-6 kDa, Carl Roth, Karlsruhe, Germany) and concentrated using 5 kDa Vivaspin^®^ Turbo 4 ultrafiltration columns (Sartorius Stedim Lab Ltd., Stonehouse, UK). Protein concentrations were determined with a NanoPhotometer^®^ (Implen, California, USA).

### Production of anti-muZP2 and anti-huZP2 antibodies

Monoclonal antibodies against murine ZP2 (amino acids 114-129; clone IE-3, rat IgG) and human ZP2 (clone H2.8, mouse IgG1) were generated using hybridoma cell lines CRL-2463^TM^ and CRL-2568^TM^ (ATCC, Manassas, Virginia, USA), respectively, as previously described (Wiesehöfer et al., 2022). Antibodies were purified from hybridoma supernatants by affinity chromatography with HiTrap™ Protein G HP columns (GE Healthcare, Little Chalfont, UK) according to the manufacturer’s instructions. The antibody solutions were further dialyzed overnight (15.9 kDa pore size; Roth, Karlsruhe, Germany) to ensure purity and pH neutrality. Concentration was achieved using 10 kDa Amicron Ultra centrifugal filters (Merck Millipore, Burlington, MA, USA; 2540 x g, 20 min) followed by sterile filtrating (0.22 µm; Millex®-GV, Merck Millipore, Burlington, MA, USA). Antibody specificity toward N-terminal muZP2^35-149^ or huZP2^39-154^ was confirmed by ELISA and Western blot analysis.

### Solid Phase Enzyme-linked Immunosorbent Assay (ELISA)

Specificity of the generated anti-muZP2 and anti-huZP2 antibodies for the N-terminal muZP2^35-149^ or huZP2^39-154^ proteins was assessed by solid phase ELISA. In brief, 100 µl cell culture supernatant containing recombinant muZP2^35-149^ or huZP2^39-154^ was coated onto 96-well plates (Thermo Fisher Scientific, Oberhausen, Germany) overnight at 4°C. Plates were blocked with PBS containing 1% bovine serum albumin (BSA; Roth, Karlsruhe, Germany) for 2 h at RT, washed twice with PBS and then incubated with 17 µg/ml anti-muZP2 antibody (IE-3, CRL-2463™; ATCC^®^) or 10 µg/ml anti-huZP2 antibody (H2.8, CRL-2568™; ATCC^®^) diluted in PBS containing 0.5% BSA for 2 h at RT or 12 h at 4°C. After three washing steps, wells were incubated with HRP-conjugated goat anti-rat (for muZP2) or goat anti-mouse (for huZP2) secondary antibodies for 90-120 min. Following four washes, 100 µl TMB substrate (Biotrend Chemikalien GmbH, Cologne, Germany) was added, and the reaction was stopped after 15 min with 100 µl of 0.2 M H_2_SO_4_ (Carl Roth, Karlsruhe, Germany). Absorbance at 450 nm (reference 620 nm) was measured in a microplate reader (Tecan, Männedorf, Switzerland). Samples were measured in duplicates. Isotype controls (rat anti-murine IgG2a and mouse anti-human IgG1) were included. Non-specific binding of secondary antibodies was excluded. All antibody solutions were diluted in PBS containing 0.5% BSA.

### Protein isolation

#### Validation of N-terminal ZP2 peptide production

Cell culture supernatants or 1 µg of the purified recombinant peptide were mixed with Laemmli buffer (62.5 mM Tris, 2% SDS, 25% glycerol, 0.01% bromophenol blue, 5% β-mercaptoethanol).

#### Protein isolation of murine and human spermatozoa

Isolation was performed as previously described (Wennemuth *et al*, 2000):In short, non-capacitated and capacitated sperm were washed twice with HS-buffer by centrifugation (300 x g, 5 min) and pellets were resuspended in HS (15 µl per 0.5 x 10^6^ spermatozoa), then mixed with 2x Laemmli buffer (in mM: 278 SDS, 120 Tris-HCl pH 8.5, 0.1 bromophenol blue, 2740 Glycerin) in a 1:1 ratio. Samples were boiled for 5 min at 95°C, centrifuged (300 x g, 5 min, 4°C), then supplemented with 5% ß-mercapthoethanol (Sigma-Aldrich, Darmstadt, Germany) and boiled for another 5 min at 95°C. Protein were stored at –80°C.

#### Digestion and heat-inactivation of muZP2^35-149^

MuZP2^35-149^ was enzymatically digested by the use of Mag-Trypsin (TPCK-trypsin immobilized on magnetic beads; Cat. No. 635646; Takara Bio, Saint-Germain-en-Laye France) according to the manufacturer’s specifications. In short: 1 mg protein was diluted in digestion buffer (0.1 M NaHCO3, pH 8.3, 6-8 M urea) and digested by incubating for 15 min at 95°C. Urea concentration was reduced to >1 M by one washing step (0.1 M NaHCO3, pH 8.3). Protein mixture was added to a Mag-Trypsin suspension and incubated over night at 37°C with vigorous mixing. The digested peptide mixture was collected using a magnetic separator and stored at –80°C.

Heat-inactivation of muZP2^35-149^ was performed by boiling muZP2^35-149^ at 95°C for 5 min. The heat-inactivated protein was used in the relevant experiments without further processing. Successful digestion and heat-inactivation was proved by SDS-PAGE and silver staining.

### Immunoblot

Samples (1 µg purified protein or 0.5 x 10^6^ sperm/lane) were subjected to electrophoresis on 8-16% TGX stain-free gels (Bio-Rad, Feldkirchen, Germany) and transferred to 0.2 µm PVDF membranes). Membranes were blocked with 5% BSA/PBS and incubated overnight with primary antibodies as follows: anti-muZP2 monoclonal rat antibody (IE-3, CRL-2463™; ATCC^®^) in a concentration of 1:500; anti-huZP2 monoclonal mouse antibody (H2.8, CRL-2568™; ATCC^®^) in a concentration of 1:1000; alpha-Tubulin (clone B-5-1-2, 1:1,000, Merck Millipore Ltd.); CatSper1 (clone 6D3, 1:1,100, SigmaMillipore) (Hwang *et al*, 2025).

HRP conjugated goat anti-rat (clone 31470, Thermo Fisher Scientific, Oberhausen, Germany) or HRP conjugated goat anti-mouse (Jackson Immuno Research, London, England) secondary antibodies were used for detection. Protein bands were visualized using Clarity™ Western ECL Substrate and a ChemiDoc™ Touch Imaging System (Bio-Rad, Düsseldorf, Germany).

### Protein Multimer prediction

The mouse CatSper canopy protein sequences were adopted from the PDB structure (7EEB) as the binding partners for the murine or human ZP2 protein sequences (Lin *et al*, 2021). Homolog sequences of human CatSper canopy were predicted by comparison with 7EEB (CTSRE_HUMAN, Q5SY80; CTSRD_HUMAN, Q86XM0; CTSRB_HUMAN, Q9H7T0; CTSRG_HUMAN, Q6ZRH7). The prediction of the protein multimer was performed using AlphaFold 3 (Abramson *et al*, 2024). The predicted multimers are aligned with the entire CatSpermasome structure from 7EEB using UCSF ChimeraX (Meng *et al*, 2023).

### Amino acid alignment

Amino acid alignments were analyzed with the using BLAST algorithm(Altschul *et al*, 1997; Altschul *et al*, 2005).

### Assessment of acrosomal exocytosis

Spermatozoa from SU9-DsRed/Acr3-EGFP mice were prepared as described and incubated with either 20 µM ionophore A23187 (control) or 200 ng/ml muZP2^35-149^. Subsequently, 25-30 µl of the sperm suspension was placed onto Superfrost^TM^ Plus slides (25×75×1 mm; R. Langenbrinck GmbH, Emmendingen, Germany) for 30 min at RT to facilitate attachment, followed by fixation in methanol (15 min, RT). Acrosomal exocytosis was assessed using a Ti-U Eclipse microscope (Nikon, Düsseldorf, Germany) equipped with a 40x objective.

Additionally, acrosomal status of fixed wildtype spermatozoa was analyzed by first treating them with either 20 µM ionophore A23187 (control) or 200 ng/ml muZP2^35-14^ and then staining them with fluorescein isothiocyanate-conjugated (FITC)-conjugated pisum sativum agglutinin (PSA; 12.5 µg/ml; Sigma-Aldrich, St. Louis, MO, USA). Labeled spermatozoa were mounted in Vectashield^®^ antifade mounting medium containing DAPI (H-2000; Vector Laboratories, Susteren, Netherlands) to prevent photobleaching. For each condition, 150 cells per slide were evaluated at 400x magnification.

### Tissue preparation, immunofluorescence and immunohistological staining

Tissues were fixed using 4% formalin and embedded into paraffin. FFPE tissues were cut into 4 µm sections using a microtome, deparaffinized, rehydrated and stained.

#### Immunofluorescence staining

After deparaffinization, sections were pre-treated with Vectastain Elite ABC Kit (Vector laboratories, Newark, CA, USA) according to manufacturer’s instructions to reduce tissue autofluorescence. Antigen retrieval was performed by heating in citrate buffer (11 mM citric acid, pH 6.0) for 30 min. Sections were permeabilized with 0.1% Triton X-100 in PBS for 10 min and blocked with PBS containing 1% BSA and 10% goat serum for 60 min. Sections were incubated with the anti-huZP2 (H2.8; 1:20) or anti-muZP2 (IE-3, 1:50) antibodies overnight at 4°C. After washing, the sections were incubated with secondary antibody Alexa Fluor^®^ 488 AffiniPure^®^ F(ab’)2 Fragment Goat anti-Mouse IgG (H+L) or Alexa Fluor^®^ 488 AffiniPure^®^ Goat Anti-Rat IgG (H+L) (1:200, Jackson ImmunoResearch, Biozol, Hamburg, Germany) solution containing DAPI (1:200) for 60 min. Sections were washed, mounted with Fluoromount (Southern Biotech, Birmingham, AL, USA) and visualized using a Ti2-E microscope (Nikon, Düsseldorf, Germany) with 40x objective.

#### Immunohistological staining

Following deparaffinization, antigen retrieval was performed in citrate buffer (pH 6.0) for 30 min at 95°C. Sections were permeabilized with 0.1% Trinton X-100 (in PBS) for 10 min, washed three times with PBS, and blocked with 1% BSA/PBS. Sections were incubated with anti-huZP2 (H2.8, 1:100) or anti-muZP2 (IE-3, 1:400) antibodies in 1%BSA/PBS overnight at 4°C. After two PBS washes, slides were incubated for 1h at RT with rabbit anti-mouse IgG Fc (Biotin) (Thermo Fisher Scientific, Oberhausen, Germany) or rabbit anti-rat IgG (Biotin) (DAKO, Hamburg, Germany) diluted 1:500 in PBS. Slides were washed twice, and signals were amplified using a Streptavidin-Biotin complex (Thermo Fisher Scientific, Oberhausen, Germany). 3,3’-diaminobenzidine (DAB, Sigma Aldrich, Darmstadt, Germany) and H_2_O_2_ solution were used for detection. Sections were counterstained with Instant hematoxylin (Thermo Fisher Scientific, Oberhausen, Germany), dehydrated and mounted using Xylene Substitute mounting medium (Thermo Fisher Scientific, Oberhausen, Germany).

### Dye loading and photometry

For Ca^2+^-Imaging sperm were incubated in HS buffer containing 5 µM Fluo-4 AM (Thermo Fisher Scientific, Oberhausen, Germany) and 10% Pluronic^®^ F-127 (Thermo Fisher Scientific) for 45 min at 37°C. After one washing step, sperm were allowed to adhere to a glass slide for 10 min. Recordings were performed with a Nikon Ti2-E microscope (Nikon, Düsseldorf, Germany) equipped with a CFI P-Apo VC 60X water immersion objective. Fluo-4 AM was excited using a Lumencor SpectraX light source (EX 482/35; Lumencor, Beaverton, USA) at 1-5 % power to minimize photobleaching, and fluorescence emission (496 nm and 576 nm) was detected by an Orca Fusion BT camera (Hamamatsu, Hamamatsu City, Japan). Cells were continuously perfused at RT with a micropipette as previously described (Wennemuth *et al*., 2000) either with HS (first 10 s) or with 200 ng/ml muZP2^35-149^ in HS (75 s). Imaging analysis was performed using NIS-Elements software (Vers. 6.0; Nikon, Düsseldorf, Germany).

For photometric measurement of sperm suspensions, dye loading was performed as described above. The cell suspension was transferred to 96-well-plates (Greiner Bio-One GmbH, Frickenhausen, Germany), and the basal fluorescence was recorded for 20 s using a ClarioStar Plus plate reader (BMG Labtech, Ortenberg, Germany) equipped with an automated injection system. Compounds were added in a 1:1 ratio, and emission intensity was monitored for additional 37 s. Ionophore A23187 (5 µM final concentration) was used as a positive control. Ca^2+^ signal (Δ*F*/*F* _0_) 20 s after stimulation were calculated as (F – F0)/F0, where F represents fluorescence after stimulation and F0 the baseline fluorescence immediately prior to stimulation, as indicated in the figure legends.

### Digital Holographic imaging for 4D sperm motility analysis

Digital holographic microscopy was used for 4D motility analysis of murine and human spermatozoa as previously described (Muschol *et al*, 2018; Wiesehofer *et al*., 2022). Briefly, measurements were performed using a DHM™ T-1000 digital holographic microscope (Lyncée Tec SA, Lausanne, Switzerland) operating in an off-axis transmission mode, equipped with a 666 nm laser diode, a 40x/0.6 NA objective, and a Basler aca1920-155um CCD camera (Basler AG, Ahrensburg, Germany). Experiments were conducted at 37°C in 100 µm deep chamber slides (Leja products B.V., Nieuw-Vennep, Netherlands) at a recording rate of 100 fps. Holographic images were processed offline using Koala software (Vers. 6; Lyncée Tec SA) and open-source Spyder (Python v3.6.9). Motility parameters like curvilinear velocity (VCL) and the 2D amplitude of lateral head displacement (ALH) were calculated from reconstructed X, Y and Z coordinates. Flagellar movements were analyzed by tracing individual frames in stacks of XY projections (8-bit TIFF, 100 fps, 10 frame segments) using a custom macro in Igor Pro™ Vers. 6.36 (Wavemetrics, Lake Oswego, OR, USA). Brightness and contrast of reconstructed XY projections were adjusted with ImageJ V1.50i (NIH). Z-plane coordinates were determined using custom scripts in Spyder; conversion from pixel to micrometers was performed using a P/U value of 7.49, and smoothing of Z-plane data applied a seventh-order polynomial in Igor Pro^TM^. For 4D motility analysis, non-capacitated or capacitated human sperm were washed twice, re-suspended to a final concentration of 1x10^6^ sperm/ml and analyzed with the DHM. For experiments, 30 µl of sperm suspension (w/o beads as control, or in presence of either unloaded or huZP2^39-154^ –loaded beads or soluble huZP2^39-154^) was transferred to the chamber slide. Only free-swimming spermatozoa that had detached from the beads during transfer to the measurement chamber were analyzed.

### Statistical analysis

Statistical analysis was performed with GraphPad Prism (Vers.10, Statcon GmbH, Witzenhausen, Germany). One-way and two-way analyses of variance (ANOVA) were applied to assess differences in mean or median values, as appropriate. A p-value < 0.05 was considered statistically significant. Data are presented as medians and interquartile ranges or as mean ± SEM as indicated in the figure legends.

## Results

### Recombinant ZP2 N-terminal fragments from mouse and human induce a robust Ca²⁺ influx in capacitated murine and human spermatozoa

To investigate the function of the ZP2 N-terminal fragment, we first validate species-specific monoclonal antibodies against murine and human ZP2 (i.e., muZP2 and huZP2, respectively). Immunohistochemistry (IHC) and immunofluorescence (IF) analyses confirmed that each antibody specifically stained the zona pellucida surrounding oocytes in their respective ovarian tissues (**Fig. 1**, A-B), with further validation provided by ELISA (**Supplementary Fig. S1A**).

**Figure 1:**
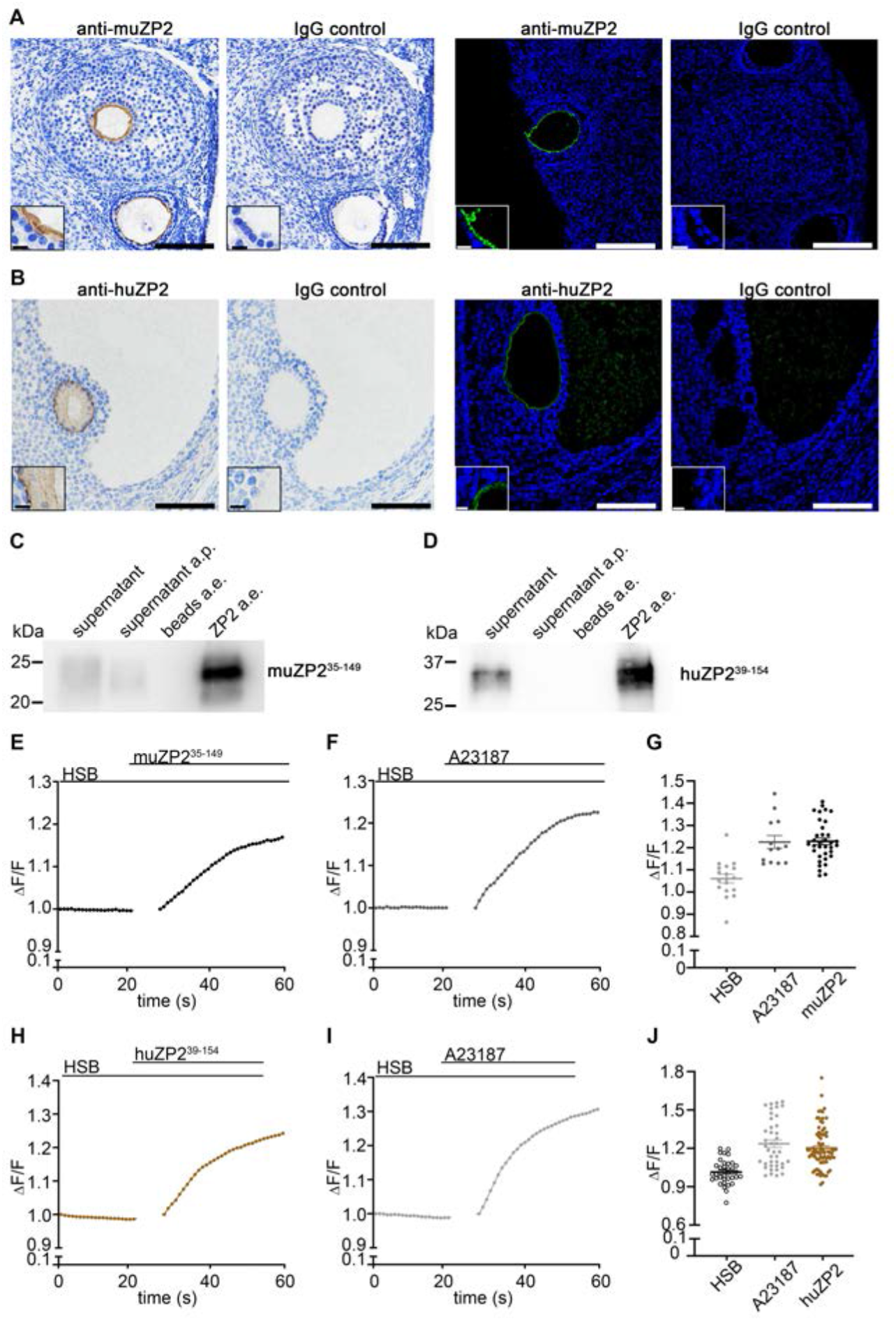
Recombinant N-terminal ZP2 fragments from mouse and human are correctly expressed and induce a Ca^2+^ signal in murine and human spermatozoa. (A-B) Validation of species-specific ZP2 antibodies. Immunohistochemistry (left) and immunofluorescence staining (right) show specific staining of the zona pellucida in mouse (A) and human () ovarian tissue. Scale bar: 100 μm, insert: 10 μm. (C-D) Production of recombinant ZP2 fragments. Western blots confirm the successful purification of N-terminal murine ZP2^35–149^ (C) and human ZP2^39–154^ (D) from HEK293T cells. Lanes 1–3 show 20 μl sample from various purification steps [a.p., after purification; a.e., after IMAC elution]; lane 4 shows 1 μg of the final purified protein. (E-J) Photometry reveals an increase in [Ca^2+^]_i_ in Fluo-4–loaded murine (E-G) and human sperm (H-J) following stimulation with either 200 ng/ml muZP2^35-149^ (E+G) or huZP2^38-154^ (H+J), with a maximal response close to that induced by the ionophore A23187 (F+I). (G+J) shows the mean values from 34 experiments for muZP2^35-149^ (G) and from 77 experiments for huZP2^38-154^ (J) and the mean values after A23187 stimulus of 12 experiments with murine sperm (G) and of 40 experiments with human sperm, each with at least 6.25 Mio cells per ml.

Next, we expressed and purified recombinant N-terminal fragments of muZP2 (amino acids 35–149; muZP2^35-149^) and huZP2 (amino acids 39–154; huZP2^39-154^). Western blot analysis confirmed the successful production of both proteins at their expected sizes (∼22 kDa; ∼30 kDa, respectively) (**Fig. 1**, C-D). These results demonstrate that our recombinant N-terminal ZP2 fragments are correctly produced.

To quantify the effect of ZP2 on sperm Ca^2+^ signaling, we monitored intracellular Ca^2+^ in Fluo-4-loaded non-capacitated and capacitated murine and human spermatozoa (**Fig. 1**, E-J; **Supplementary Fig. S1,** B-D, **Supplementary Fig. S2A**) using a plate reader equipped with an automated injection system. Stimulation with muZP2^35-149^ (final concentration of 200 ng/ml, pH 7.4) triggered an immediate and significant increase in intracellular Ca^2+^ ([Ca^2+^]_i_) compared to a physiological buffer (HSB) control (ΔF/F: 1.06 ± 0.1) in capacitated but not in non-capacitated spermatozoa (**Fig. 1**, E and G, **Supplementary Fig. S1**, B-E). Strikingly, the magnitude of this ZP2-induced response (ΔF/F: 1.23 ±0.1) in capacitated sperm was indistinguishable from the maximal influx triggered by a saturating concentration of the Ca^2+^ ionophore A23187 at 20 μM (**Fig. 1**, F-G, **Supplementary Fig. S1**, E-F).

To determine if the muZP2-induced [Ca^2+^]_i_ is a conserved mechanism in humans, we tested the effect of the recombinant human ZP2 N-terminus (huZP2^39-154^) on non-capacitated (**Supplementary Fig. S2A**) and capacitated human sperm (**Fig. 1**, H+J). Stimulation with 200 ng/ml recombinant huZP2^39-154^ triggered a rapid and robust rise in [Ca²⁺]_i_ (ΔF/F: 1.20±0.2) only in capacitated sperm, comparable in magnitude to the maximal response elicited by the ionophore A23187 (ΔF/F: 1.2±0.2) (**Fig. 1**, H-J; **Supplementary Fig. S2B**).

This demonstrates that the ZP2 N-terminus is sufficient to elicit a robust Ca^2+^ response in capacitated murine and human sperm.

### The ZP2-evoked, tail-to-head Ca^2+^ signal requires the CatSper channel in mice

We used time-lapse, live-cell imaging of individual Fluo-4-labelled mouse spermatozoa to determine the spatio-temporal dynamics of the Ca²⁺ response after local perfusion with muZP2^35-149^ to investigate CatSper-dependent Ca²⁺ influx. Consistent with the past studies that showed the CatSper-dependent tail-to-head propagation of the Ca²⁺ signal in mouse sperm (Olson *et al*, 2010; Xia *et al*, 2007), stimulation with the ZP2 fragments triggered a rapid rise in [Ca^2+^]_i_ that was noted in the midpiece approximately 5 seconds after application, before propagating to the head region with a slight delay (**Fig. 2**, A-B). The signal was transient, with [Ca^2+^]_i_ returning towards baseline levels upon washout of the ZP2 fragment (**Fig. 2B**).

**Figure 2:**
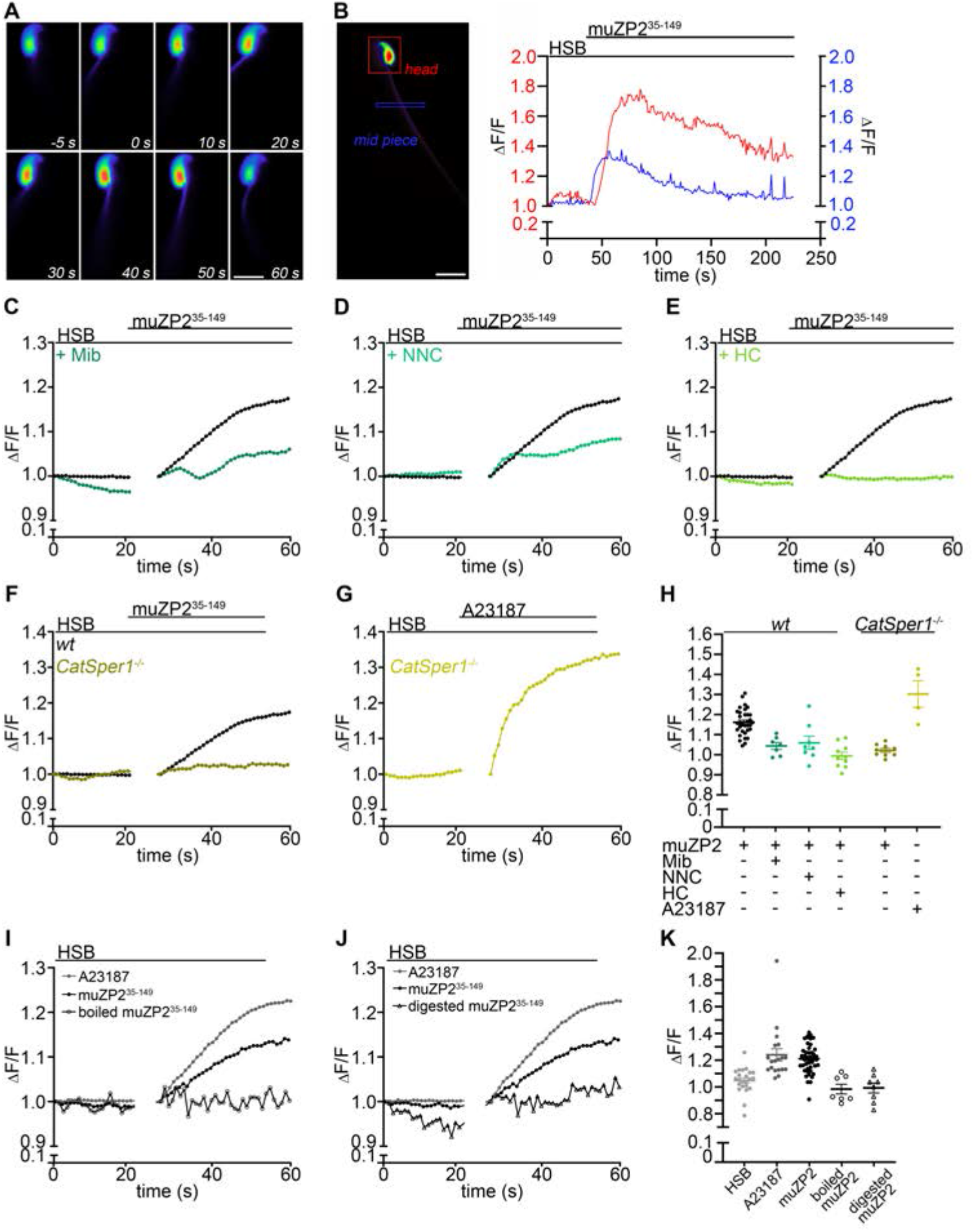
The ZP2-evoked, tail-to-head Ca^2+^ signal requires the CatSper channel in mice. (A-B) single-cell photometry of murine spermatozoa stimulated with 200 ng/mL muZP2^35-149^ for 30 s (starting at time 0 s) shows [Ca^2+^]_i_ increase in the head regions and mid piece. (B) Local recordings over time of the muZP2^35-149^-induced [Ca^2+^]_i_ signal show an initial [Ca^2+^]_i_ rise in the midpiece (blue), followed by an increase in the head region (red). Scale bar: 10 μm. (C-E) demonstrate the muZP2^35-149^-induced increase in [Ca^2+^]_i_ in the presence of different CatSper inhibitors (green) compared to the control without inhibitors (black). Mibefradil (Mib), NNC 55-0396 (NNC), and HC-056456 (HC) each reduced the response to muZP2^35-149^. No increase in [Ca^2+^]_i_ in response to muZP2^35-149^ was observed in spermatozoa from *CatSper1^−/−^* mice (F), whereas the response to A23187 stimulation remained unaffected (G). (H) statistical analysis of muZP2^35-149^ induced [Ca^2+^]_i_ response. (I-K) Photometry analyses reveal an absence of [Ca^2+^]_i_ in Fluo-4–loaded murine sperm following stimulation either with 200 ng/ml boiled (I) or digested (J) muZP2^35-149^. (K) statistical analysis of photometry analyses. N_experiments_≥ 3 experiments each with at least 6.25 Mio cells per ml.

To further test whether the molecular basis for this ZP2-induced tail-to-head Ca^2+^ signal, we used both pharmacological and genetic approaches. First, we pre-incubated sperm with three distinct CatSper inhibitors: Mibefradil (Mib), NNC 55-0396 (NNC), and the highly selective HC-056456 (HC). All three compounds abolished the ZP2-evoked Ca^2+^ increase in a dose-dependent manner (**Fig. 2**, C-E, **Supplementary Fig. S1**, G-H). Notably, 10 μM concentrations of Mib or NNC reduced the Ca^2+^ influx by over 90%, while the highly selective inhibitor HC achieved near-complete blockade at just 5 μM (muZP2^35-149^= ΔF/F: 1.23±0.1; HC= ΔF/F: 1.0 ± 0.1) (**Fig. 2**, C-E, **Supplementary Fig. S1**, G-H). When the assay was performed on the sperm from CatSper1 knockout mice in parallel, these cells failed to produce a Ca^2+^ signal in response to muZP2^35-149^, yet they exhibited a robust, wild-type-level response to the Ca^2+^ ionophore A23187 (wild-type= ΔF/F: 1.23±0.1; *CatSper1^-/-^=* ΔF/F: 1.34±0.1), confirming their viability (**Fig. 2**, F-H; **Supplementary Fig. S1I**).

To confirm that the Ca²⁺ response depends on the intact, correctly folded ZP2 fragment rather than on a non-specific effect of the ZP2 preparation, we stimulated capacitated murine sperm with muZP2^35-145^ that had been either heat-denatured or proteolytically digested. Neither treatment elicited a detectable [Ca²⁺]ᵢ increase (Fig. 2, I–K), demonstrating that the structural integrity of the N-terminal ZP2 fragment and ruling out a contribution of buffer components, contaminants, or degradation products (**Fig. 2**, I-K).

Taken together, these pharmacological and genetic data conclusively demonstrate that the ZP2-induced increases in intracellular Ca²⁺, with tail to head propagation, is entirely dependent on the structural integrity of muZP2^35-145^ and a functional CatSper channel.

### The N-terminal fragment of human ZP2 functions as a CatSper activator, similar to progesterone

To determine if the ZP2-CatSper signaling pathway is a conserved mechanism in humans, we tested the effect of huZP2^39-154^ on capacitated human sperm by pre-treatment with three distinct CatSper inhibitors at 5 and 10 µM (Mib, NNC, and HC) (**Fig. 3**, A-C and F; **Supplementary Fig. S2**, C-E). The huZP2^39-154^ signal was CatSper-dependent, as it was completely abrogated by pre-treatment with the three distinct CatSper inhibitors.

**Figure 3:**
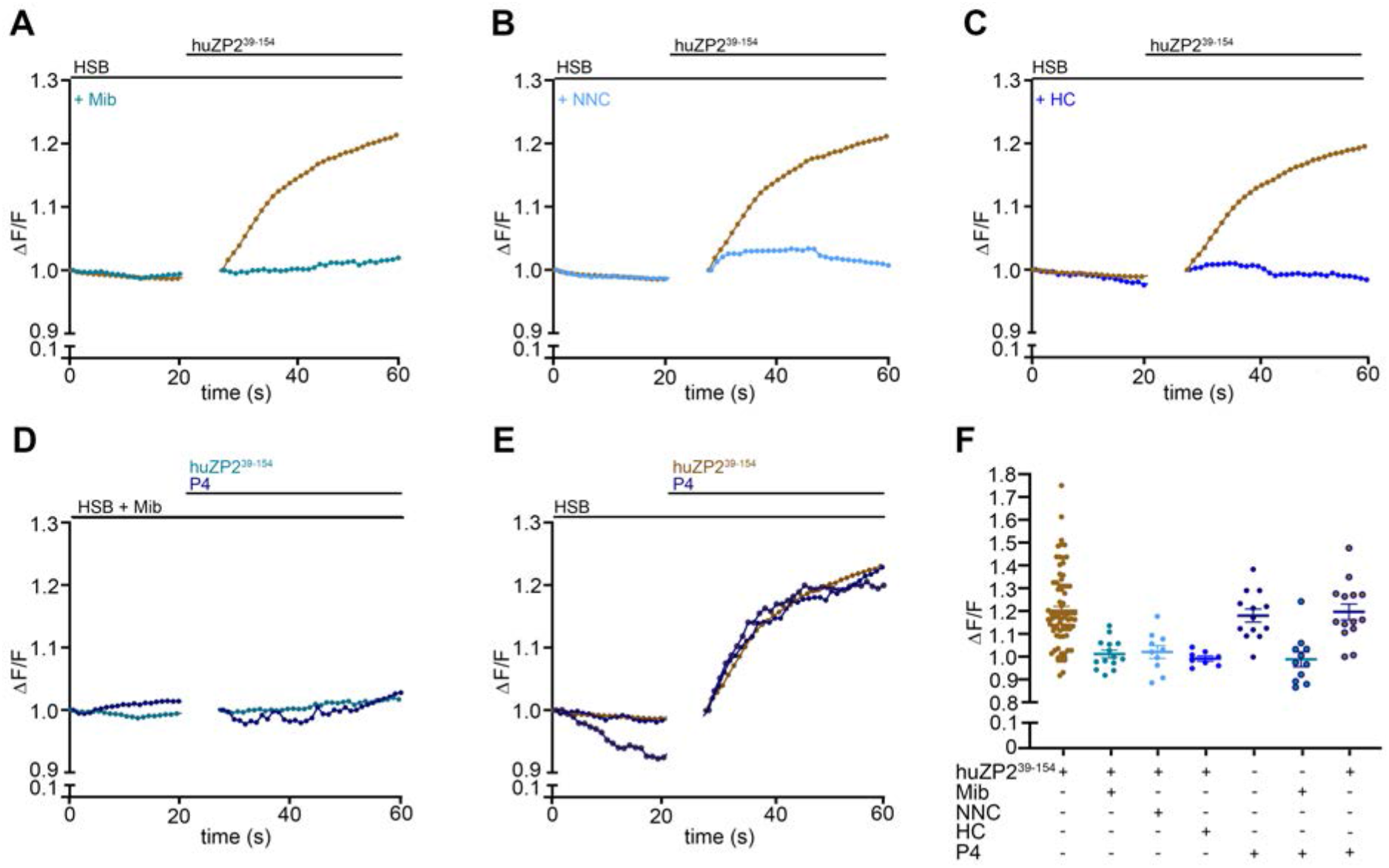
The human ZP2 N-terminus is a potent agonist of the human CatSper channel. (A-F) Photometric analysis of Fluo-4–loaded human spermatozoa demonstrates a robust increase in [Ca²⁺]_i_ following stimulation with 200 ng/ml huZP2^38-154^ (orange). [Ca²⁺]_i_ increase induced by huZP2^38-154^ in the presence of CatSper inhibitors (various blue shades): 5 µM Mibefradil (Mib), NNC 55-0396 (NNC), or HC-056456 (HC), each markedly reducing the Ca²⁺ influx compared to control. (D) Co-application of huZP2^38-154^ or P4 with 5 µM Mib results in reduced Ca²⁺ entry. (E) Comparison of the [Ca²⁺]_i_ increase elicited by huZP2^38-154^, by 500 nM progesterone (P4) or by simultaneous addition of huZP2^38-154^ and P4 shows similar amplitudes. (F) Statistical analysis of the Ca²⁺ experiments shown in A-E. N_experiments_≥ 8 experiments each with at least 6.25 Mio cells per ml.

To benchmark this activity, we compared the huZP2-evoked signal to that of progesterone (P4), the best-characterized and potent agonist of human CatSper (Lishko *et al*, 2011b; Strünker *et al*, 2011). Remarkably, huZP2^39-154^ and P4, induced Ca²⁺ signals of nearly identical amplitude (ΔF/F: 1.20±0.2 and 1.18±0.1) (**Fig. 3**, E-F; **Supplementary Fig. S2F**). Simultaneous application of huZP2^39-154^ and P4 shows no additive effect (ΔF/F: 1.20±0.1). As all tested agonists evoked responses close to the upper range of the Fluo-4 signal, this lack of additivity most likely reflects saturation of the calcium indicator rather than proof of a shared binding site, and is consistent with a common, saturable CatSper-dependent pathway. Furthermore, both responses showed comparable sensitivity to CatSper inhibition by 5 µM Mib (huZP239-154= ΔF/F: 1.01±0.1, P4= 1.0±0.1) (**Fig. 3**, D and F; **Supplementary Fig. S2G**). This pharmacological profile provides evidence that the N-terminal fragment of human ZP2 acts as a potent physiological agonist for the human CatSper channel, establishing a key molecular link between the human egg and sperm.

### The human ZP2 N-terminus induces hyperactivation-like motility and acrosomal exocytosis

To determine if the huZP2-evoked Ca^2+^ signal translates to functional changes in motility, we used digital holographic microscopy (DHM) (Wiesehofer *et al*., 2022) to analyze 4D sperm movement of human capacitated sperm after contact to huZP2^38-154^ diluted in capacitating buffer (= soluble huZP2^38-154^) or to huZP2-loaded beads (**Fig. 4, Supplementary Fig. S3**). While control beads had no effect, beads coated with huZP2^38-154^ (**Supplementary Fig. S3**) and free huZP2^38-154^ (**Fig. 4**) induced significant changes characteristic of hyperactivation in capacitated sperm. Specifically, huZP2 stimulation led to a significant increase in the amplitude of the flagellar beat (XY-excursion), and the overall curvilinear velocity (VCL) of human spermatozoa (**Fig. 4**, A *first-third figures* and B *left figure*, **Supplementary Fig. S3**, A-C). These parameters collectively describe a faster, more powerful, and asymmetric flagellar beat. Interestingly, other parameters like out-of-plane flagellar motion (Z-excursion) and lateral head displacement (ALH) were largely unaffected, suggesting a specific modulation of the propulsion machinery of sperm (**Fig. 4**, A *first-second and forth figure* and B, *right*; **Supplementary Fig. S3B**, *right*). Collectively, these data demonstrate that specific interaction of the ZP2 N-terminal fragment is sufficient to switch human sperm into a hyperactivated-like state, preparing it to penetrate the oocyte vestments.

**Figure 4:**
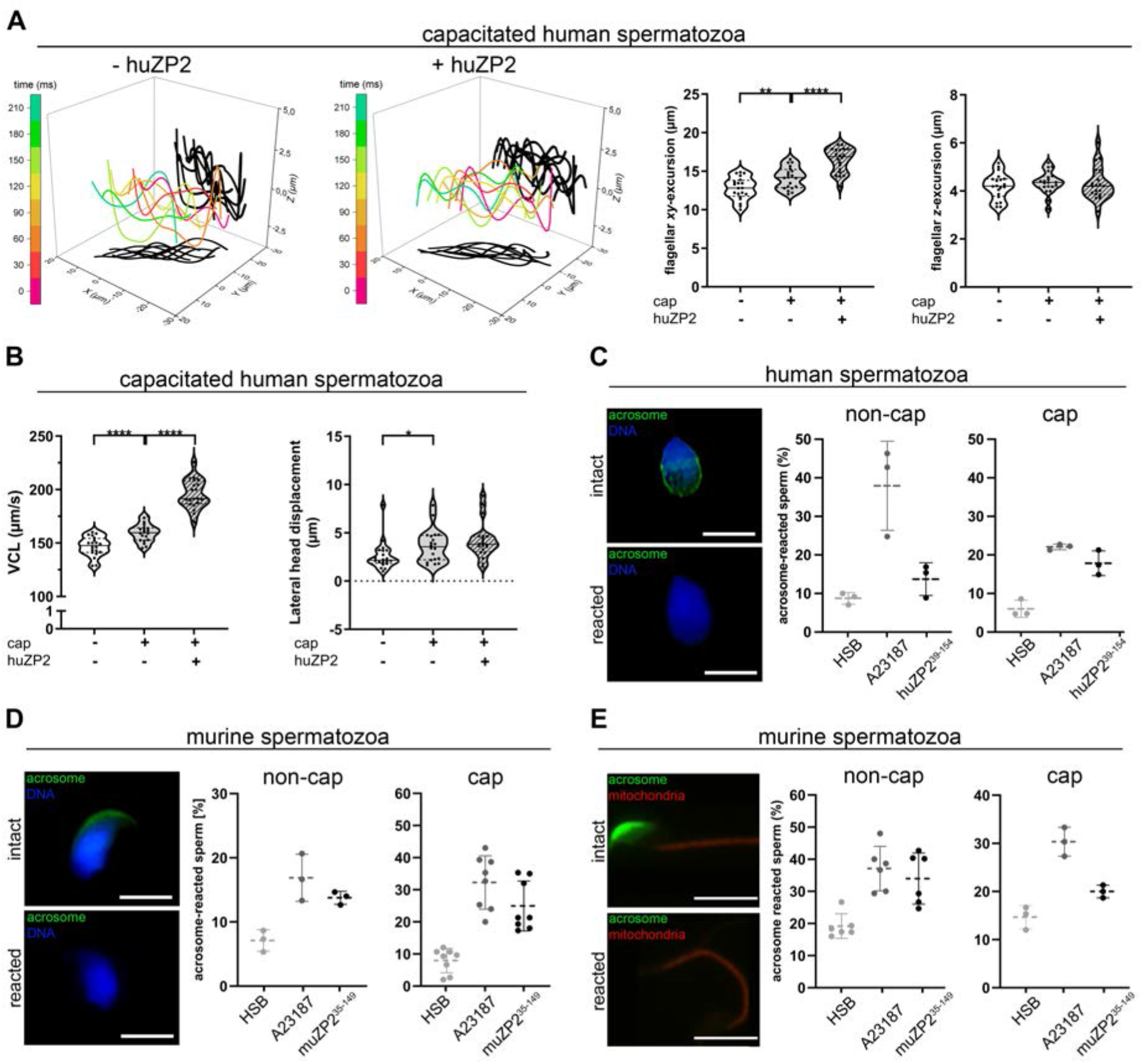
HuZP239^-154^ increases speed, flagellar XY-excursion and induces acrosomal exocytosis in human spermatozoa. (A-B) 4D motility analysis by digital holographic microscopy of non-capacitated and capacitated human sperm. (A) The flagellar waveform of capacitated (first) and capacitated human sperm after contact to huZP2^39-154^ (second) is shown for one beat cycle. The 3D flagellar excursions at different time points (0, 40, …, 210 ms) are color-coded, with their projections onto the XY– and XZ-planes shown in black. Statistical analysis of flagellar XY– (A, third), Z-excursions (A, forth), the curvilinear velocity (VCL) (B, left) and lateral head displacement (B, right). The analysis shows that the exposure to huZP2^39-154^ (=huZP2) led to a significant increase in the flagellar XY-excursion and the VCL. (C-E) Evaluation of the acrosomal reaction status of human (C) and murine spermatozoa (D-E) following huZP2^38-154^ or muZP2^35-149^ treatment (200 ng/ml) compared to A23187 treatment using PSA staining (C-D) and Su9-DsRed/Acr-EGFP spermatozoa (E). Scale bar: 10 μm.

The induction of AR occurs in the distal isthmus of the female genital tract (Hino *et al*, 2016; La Spina *et al*, 2016; Muro *et al*, 2016). Given that a partial cleavage of ZP2 even before fertilization was reported (Körschgen *et al*, 2017; Xiong *et al*, 2017a), we investigated whether the muZP2^35-149^ and huZP2^38-154^-induced Ca²⁺ influx is also sufficient to induce AR. Therefore, we labeled human and murine sperm with fluorescein isothiocyanate (FITC)-Pisum sativum agglutinin (PSA) (**Fig. 4**, C-D) and used additionally sperm from Su9-DsRed/Acr-EGFP transgenic mice (**Fig. 4E**) (Luo *et al*, 2013). Stimulation with huZP2^38-154^ or muZP2^35-149^ caused a significant increase in the acrosome reaction rate in the respective capacitated human or murine sperm compared to buffer controls (huZP2^38-154^: ΔAR »12%; muZP2^35-149^: ΔAR_PSA_ »17%, ΔAR_SU9/Acr3_ »17%) (**Fig. 4**, C-D). In all cases, the ZP2-induced acrosomal exocytosis rate (human sperm: 14-21%; murine sperm: 25-36%) was substantial, approaching the maximal response triggered by the Ca^2+^ ionophore A23187 (human sperm: 21-23%; murine sperm: 32-40%). This demonstrates that the ZP2-evoked Ca^2+^ signal is sufficient to trigger the acrosome reaction *in vitro* in a significant population of capacitated sperm.

### Structural modeling predicts a possible direct interaction of the ZP2 N-terminus with the CatSper channel

To determine if ZP2-CatSper functional link potentially involves direct interaction, we performed AlphaFold-Multimer with the CatSper canopy and the ZP2 N-terminal fragment (Abramson *et al*., 2024). The models predicted a potential binding interface between the ZP2 N-terminus and the extracellular “canopy” of the CatSper channel (**Fig. 5A**; **Supplementary Fig. S4A**).

**Figure 5.**
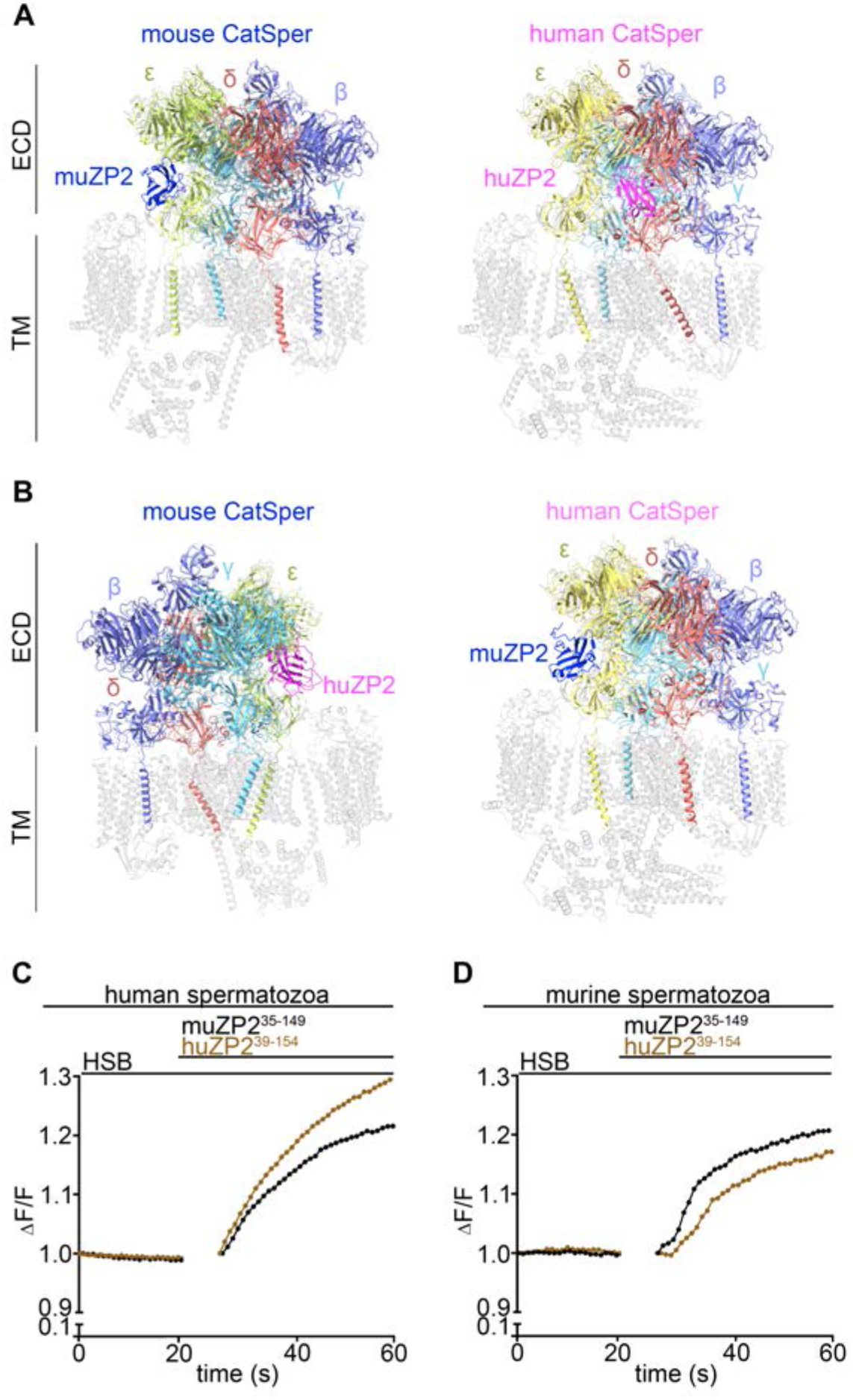
CatSper binding to recombinant human and mouse ZP2. (A) Predicted interaction of muZP2^35-149^ (blue) with murine CatSper canopy (left) and huZP2^38-154^ (magenta) with human CatSper canopy (right) by AlphaFold3. Predicted extracellular domains (ECDs) of CatSper are colored (β= light purple; γ= sky blue; δ= salmon; ε= yellow), 7EEB structure is in grey. (B) Predicted interaction of muZP2^35-149^ (blue) with human CatSper canopy (left) and huZP2^38-154^ (magenta) with murine CatSper canopy (right, rotated by 180°) by AlphaFold3. Predicted ECDs of CatSper are colored (β= light purple; γ= sky blue; δ= salmon; ε= yellow), 7EEB structure is in grey. (C-D) Comparative analysis of [Ca²⁺]_i_ responses to huZP2^39-154^ (orange) and muZP2^35-149^ (black) in human (C) and murine (D) sperm indicating that both huZP2^39-154^ (orange) and muZP2^35-149^ (black) induce a [Ca²⁺]_i_ increase independent of species, with the magnitude of influx being higher within species-matched contexts. N_experiments_≥ 7 experiments each with at least 6.25 Mio cells per ml

Crucially, the predicted ZP2-CatSper interface appeared conserved, as not only species-specific pairs (muZP2/mouse CatSper; huZP2/human CatSper) but also cross-species combinations (e.g., huZP2/mouse CatSper) (**Fig. 5B**; **Supplementary Fig.S4B**). By comparing amino acid sequences of the N-terminal muZP2^35-149^ and huZP2^39-154^, a sequence similarity (66%) was identified (**Supplementary Fig. S4C**). Predicted cross-reactivity was determined by using Ca^2+^ photometry (**Fig. 5**, C-D; **Supplementary Fig. S4D**). Indeed, human ZP2 (huZP2^39-154^, orange trace) triggered a Ca²⁺ response in mouse sperm, and conversely, mouse ZP2 (muZP2^35-149^, black trace) activated human sperm (**Fig. 5**, C-D; **Supplementary Fig. S4D**). As expected, the extent of the Ca^2+^ response was significantly higher when the ZP2 ligand and the sperm were from the same species. This pattern of functional conservation combined with species-specific tuning is consistent with ZP2 acting as a conserved, possibly co-evolved ligand of the CatSper pathway.

Together, these structural modeling and functional data provide a molecular pathway from egg protein to sperm ion channel activation.

Our findings establish the N-terminal domain of ZP2 as a potent activator of CatSper. Exposure of capacitated sperm to soluble N-terminal ZP2 fragment initiates a flagellum-originating Ca^2+^ influx that propagates to the head, thereby triggering a defined activation cascade. This influx drives some of the hallmark features of hyperactivated motility, such as increased flagellar beat amplitude, and curvilinear velocity, likely optimizing propulsion for penetration into the oocyte. Concomitantly, ZP2-evoked Ca^2+^ signals elicit acrosomal exocytosis at rates approaching maximal ionophore stimulation, priming sperm for membrane fusion. Taken together, these results suggest of a previously unrecognized sperm activating function of N-terminal ZP2, in addition to the well-documented role in blocking polyspermy.

## Discussion

Successful mammalian fertilization is orchestrated by tightly regulated communication between the oocyte and spermatozoon. During fertilization, the Zona Pellucida (ZP) acts as an important gatekeeper, ensuring species-specific fertilization and preventing polyspermy (Avella *et al*, 2014a; Burkart *et al*., 2012). Sperm binding to the ZP leads to an intracellular Ca^2+^ increase (Balbach *et al*, 2020b) probably through an indirect activation of the sperm’s primary Ca^2+^ channel, CatSper, to complete the final stage of fertilization. However, the specific region of the ZP responsible for the observed signaling events remains unknown. Our experiments use recombinant N-terminal ZP2 peptides rather than the intact, polymerized ZP2 embedded in the zona pellucida. Thus, the effects described in this study should be interpreted as signaling elicited by a soluble ZP2 N-terminal moiety – i.e., the portion of ZP2 that can be generated and locally enriched upon ovastacin-mediated cleavage (Xiong *et al*., 2017a) – rather than as a direct proxy for the full functional repertoire of intact ZP2 in the native matrix. Thus, our work shows that soluble N-terminal fragments of ZP2 likely serve as a conserved activator of CatSper, in addition to intracellular alkalinization (Kirichok *et al*, 2006) or progesterone binding in human spermatozoa (Smith *et al*, 2013b; Strünker *et al*., 2011), refining a molecular pathway from zona-derived material to the core machinery of sperm activation.

Our findings redefine the role of ZP2, by adding a signal transduction dimension to a protein long associated with sperm-zona interactions. We demonstrate that a specific N-terminal fragment of ZP2 is sufficient to elicit a robust CatSper-dependent Ca^2+^ influx in both capacitated mouse and human sperm, with a potency rivaling that of the canonical steroid agonist progesterone (Lishko *et al*, 2011a; Lishko *et al*, 2012; Smith *et al*, 2013a; Strunker *et al*, 2011). The absolute dependence on CatSper is confirmed by pharmacological inhibition and genetic knockout. Furthermore, our spatiotemporal imaging reveals that this signal originates in the sperm tail and propagates to the head, underlining the participation of functional CatSper channels (Olson *et al*., 2010; Xia *et al*., 2007). CatSper is activated upon depolarization of the membrane potential (V_m_) (Carlson *et al*, 2003) and alkalinization of the pH_i_ (Lishko *et al*., 2011a). It remains to be determined whether ZP2’s effect on CatSper occurs indirectly through the regulation of pH_i_ or V_m_, or through direct binding to CatSper. Our structural models predict a conserved binding interface on the extracellular “canopy” of the CatSper channel, which deserves further studies for clarification.

The functional consequence of this N-terminal ZP2-evoked Ca^2+^ signal results in the induction of hyperactivated motility in murine (Wiesehöfer *et al*, 2022) and human spermatozoa, selectively enhancing the flagellar excursion and velocity required for zona penetration (Avella *et al*., 2013; Demott & Suarez, 1992; Stauss *et al*, 1995), as shown by 4D holographic microscopy. Besides sperm motility regulation, CatSper-dependent Ca^2+^ influx leads to the induction of AR (Hino *et al*., 2016; La Spina *et al*., 2016; Muro *et al*., 2016) that occurs in the distal isthmus of the female genital tract. *In vitro*, our study using recombinant N-terminal ZP2 shows that soluble N-terminal ZP2 is sufficient to induce AR in capacitated sperm across species in line with previous studies using native, isolated ZPs or solubilized ZP proteins (Balbach *et al*., 2020a; Florman & Storey, 1982; Gupta *et al*, 2012).

Regardless the earliest initial AR induction site, these results demonstrate that soluble N-terminal ZP2 fragment can trigger Ca^2+^ influx and likely accelerate AR, similar to progesterone — a potent ligand of CatSper in humans (Smith *et al*., 2013b; Strünker *et al*., 2011) — which supports our hypothesis of a CatSper-mediated signaling pathway. In an earlier study, CatSper-induced Ca²⁺ influx in humans was most strongly associated with hyperactivation, but not, as observed in this study, with AR induction (Young *et al*., 2024). Further studies are needed to fully clarify this issue.

During fertilization, cortical granule exocytosis releases the metalloproteinase ovastacin (ASTL), which cleaves ZP2 at a conserved site in the N-terminus, preventing polyspermy (Bond & Beynon, 1995; Burkart *et al*., 2012; Nishio *et al*., 2024a). The recombinant N-terminal ZP2 used in this study differs from the ZP2 fragment cleaved by Ovastacin only in that it contains 17 fewer amino acids in murine ZP2 and 13 fewer amino acids in human ZP2. Based on the very strong activation of CatSper by our N-terminal ZP2 fragment that we observed, and the finding that partial cleavage of ZP2 by ovastacin can occur even before fertilization (Körschgen *et al*., 2017; Xiong *et al*., 2017a), we speculate that ovastacin-cleaved N-terminal ZP2 fragments could “over-activate” CatSper in trailing sperm following successful fertilization. This could lead either to a functional termination or to the additional activation of “late-arriving” sperm; both events could contribute to an additional barrier against polyspermy. This idea is compatible with the updated structural and genetic model of the polyspermy block in which ZP2 cleavage primarily hardens the zona mechanically via supramolecular crosslinking (Nishio *et al*., 2024a).

In summary, our results point to a critical, conserved molecular signaling pathway between the N-terminus of ZP2 and CatSper. These new insights fundamentally advance our understanding of gamete communication and provide a powerful platform for innovations in human reproductive health. Furthermore, we postulate that this signaling pathway could provide the development of non-hormonal contraceptives and/or serves as a basis for identifying new and previously unrecognized causes of idiopathic infertility, leading to new diagnostic and therapeutic avenues.

## Acknowledgements

We dedicate this work to the memory of Dr. Donner F. Babcock, an outstanding scientist and pioneer in the field of reproductive biology, whose insightful research and collegiality have profoundly shaped our understanding of sperm physiology and fertilization. His innovative spirit, mentorship, and generosity inspired many people in the scientific community. He will be greatly missed. This work was supported by the German Research Foundation (Deutsche Forschungsgemeinschaft; WE 2344/9-1) to G.W. and by National Institutes of Health (R01HD096745) to J.J.C. J.O. was partly supported by Lalor foundation postdoctoral fellowship.

## Contributions

CW: conceptualization, methodology, formal analysis, data curation, and writing. LS, LSu, JNO, ML, TT, and LC: formal analysis and data curation. MW: conceptualization, methodology, and data curation. JJC: conceptualization, resources, supervision, and funding acquisition. GW: conceptualization, methodology, formal analysis, data curation, writing, resources, supervision, and funding acquisition.

## Ethics declarations

All experiments involving human spermatozoa were approved by the Ethics Committee of the University of Duisburg-Essen (approval number 14-5748-BO). All ovarian tissue samples in this study were collected and preserved at the Scientific Center of Pathomorphological Research, Sumy State University (Sumy, Ukraine). The use of human resected tissue specimens was approved by the Ethics Committee of Sumy State University (Protocol No. 14/65; Approval Date: March 14, 2025).

## Notes

### Competing Interest Statement

The authors have declared no competing interest.

