## Supplemental Information for "Soluble ZP2 N-terminal fragments activate CatSper-dependent Ca^2+^ entry and regulate motility and acrosomal exocytosis in mammalian sperm"

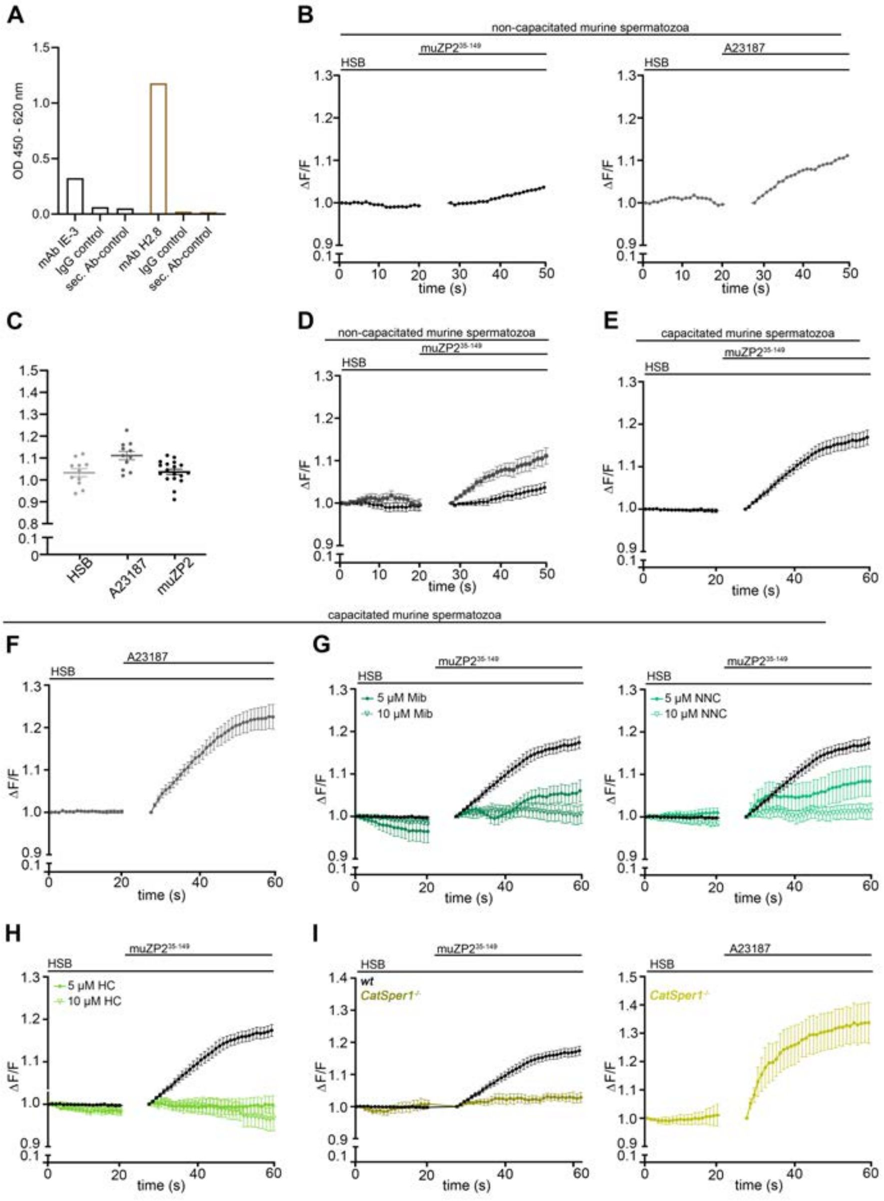


**Supplementary Fig. S1: MuZP2$^{35-149}$ induced increase in [Ca$^{2+}$]$_i$ in capacitated mouse spermatozoa can be inhibited by Mib, NNC and HC in a concentration-dependent manner**

(A) ELISA confirming specific binding of the monoclonal anti-N-terminal muZP2 antibody (IE3) to the N-terminal muZP2$^{35-149}$ peptide (lane 1) and of the monoclonal anti-N-terminal huZP2 antibody (H2.8) to the N-terminal huZP2$^{38-154}$ peptide (lane 4). IgG and secondary antibody controls (lanes 2-3 and 5-6, respectively) were negative. (B) Photometry reveals no increase in [Ca$^{2+}$]$_i$ in Fluo-4–loaded non-capacitated murine sperm following stimulation with 200 ng/ml muZP2$^{35-149}$ (left), while a maximal response induced by the ionophore A23187 was observed (right). (C) shows the mean values from 18 experiments for muZP2$^{35-149}$ and 11 experiments for A23187 stimulus, each with at least 6.25 Mio cells per ml. (D-I) Ca$^{2+}$ responses of non-capacitated (D) and capacitated spermatozoa (E-I) upon stimulation with 200 ng/ml muZP2$^{35-149}$(D-E) or with the ionophore A23187 (F). (G-I) muZP2$^{35-149}$-induced [Ca$^{2+}$]$_i$ increase was diminished in a concentration-dependent manner by CatSper channel inhibitors (green) relative to non-inhibitor controls (black). Mibefradil (Mib) (G, left), NNC 55-0396 (NNC) (G, right), HC-056456 (HC) (H). (I) No [Ca$^{2+}$]$_i$ rise was observed in *CatSper1*$^{-/-}$ -spermatozoa upon muZP2$^{35-14}$ stimulation (left), whereas the response to A23187 remained unaffected (right).

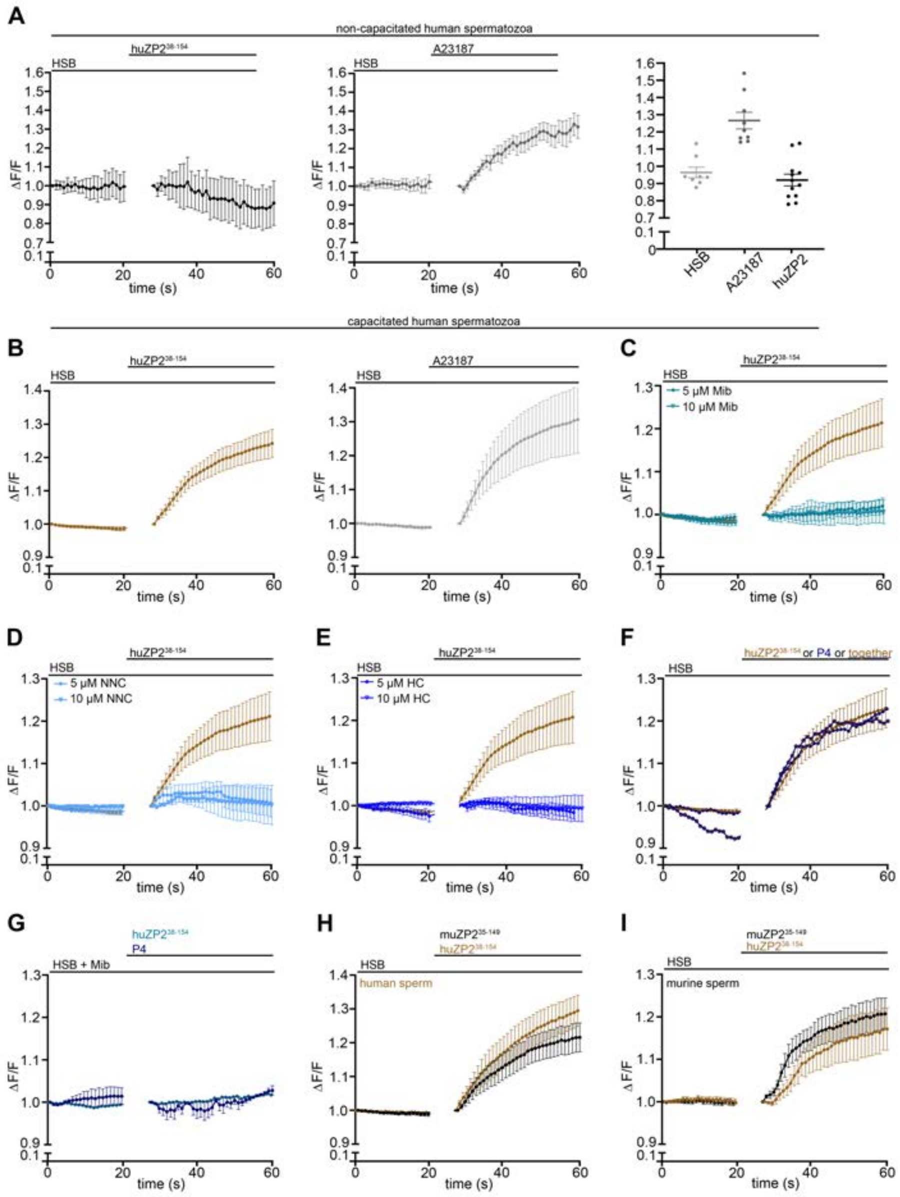


**Supplementary Fig. S2: huZP2$^{38–154}$ triggers a CatSper-dependent increase in $[Ca^{2+}]_i$ in human spermatozoa.**

(A-G) Photometry of Fluo-4–loaded non-capacitated (A) and capacitated (B-G) human sperm after stimulation either with 200 ng/ml huZP238–154 (A left-B left) or the $Ca^{2+}$ ionophore A23187 (A right-B right). $N_{experiments} \geq 8$ experiments each with at least 6.25 Mio cells per ml. (C–E) The $[Ca^{2+}]$i increase elicited by huZP238–154 in the absence (orange) or presence of CatSper inhibitors (blue shades) demonstrates a concentration-dependent effect. Mibefradil (Mib) (C) and NNC 55-0396 (NNC) (D) progressively reduce the huZP238–154–induced response, whereas HC-056456 (HC) fully suppresses the signal already at 5 µM (E). (F) The amplitude of the $[Ca^{2+}]$i increase after huZP238–154 stimulation is similar to that induced by 500 nM progesterone (P4). Simultaneous applications show no additive effect. (G) Co-application of huZP238–154 or P4 with 5 µM Mib results in no $Ca^{2+}$ entry. (H, I) Comparative analysis of the $Ca^{2+}$ responses to huZP238–154 (orange) and huZP238–154 (black) in human (H) and murine (I) sperm reveals that the response is independent of species, although the influx is generally higher when the stimulus and sperm species are matched. $N_{experiments} \geq 8$ experiments each with at least 6.25 Mio cells per ml.

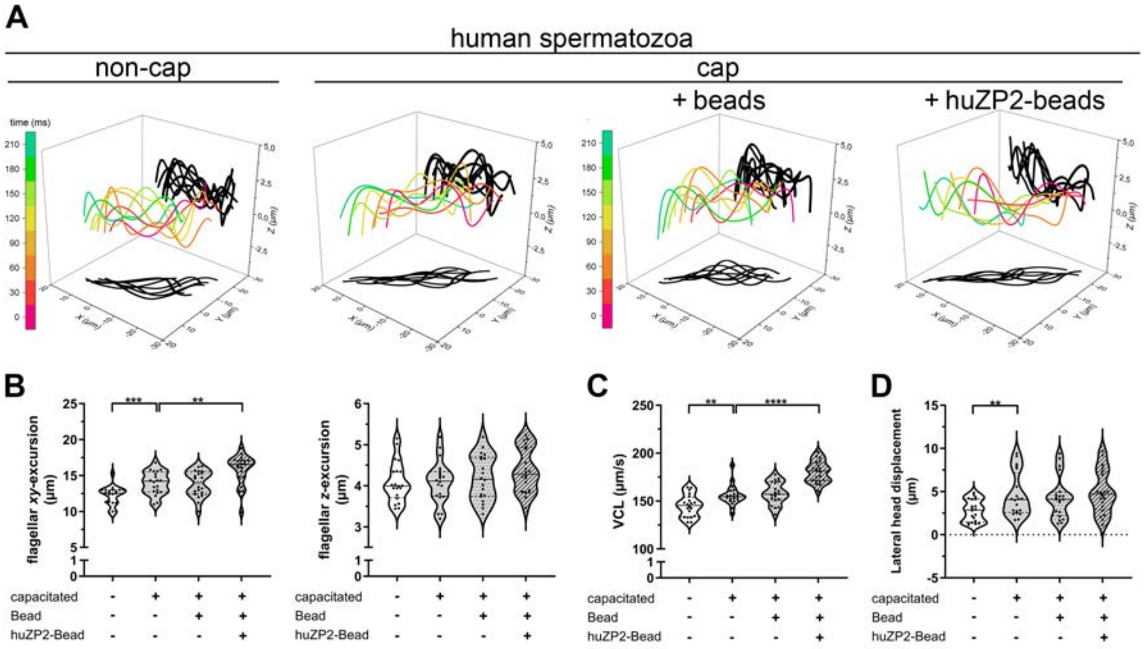


**Supplementary Fig. S3: HuZP2$^{39\text{-}154}$ increases flagellar XY-excursion and speed in human spermatozoa.**
**(**A-D) 4D motility analysis by digital holographic microscopy of non-capacitated and capacitated human sperm. (A) The flagellar waveform of non-capacitated (first), capacitated (second) or capacitated human sperm after contact to unloaded (third) or huZP2$^{39\text{-}154}$ loaded beads (forth) is shown for one beat cycle. The 3D flagellar excursions at different time points (0, 40, …, 210 ms) are color-coded, with their projections onto the XY- and XZ-planes shown in black. (B-D) Statistical analysis of flagellar XY- (B, left) and Z-excursions (B, right), the curvilinear velocity (VCL) (C) and lateral head displacement (D). The analysis shows that the exposure to huZP2$^{39\text{-}154}$-loaded beads (=huZP2-bead) led to a significant increase in the flagellar XY-excursion and the VCL.

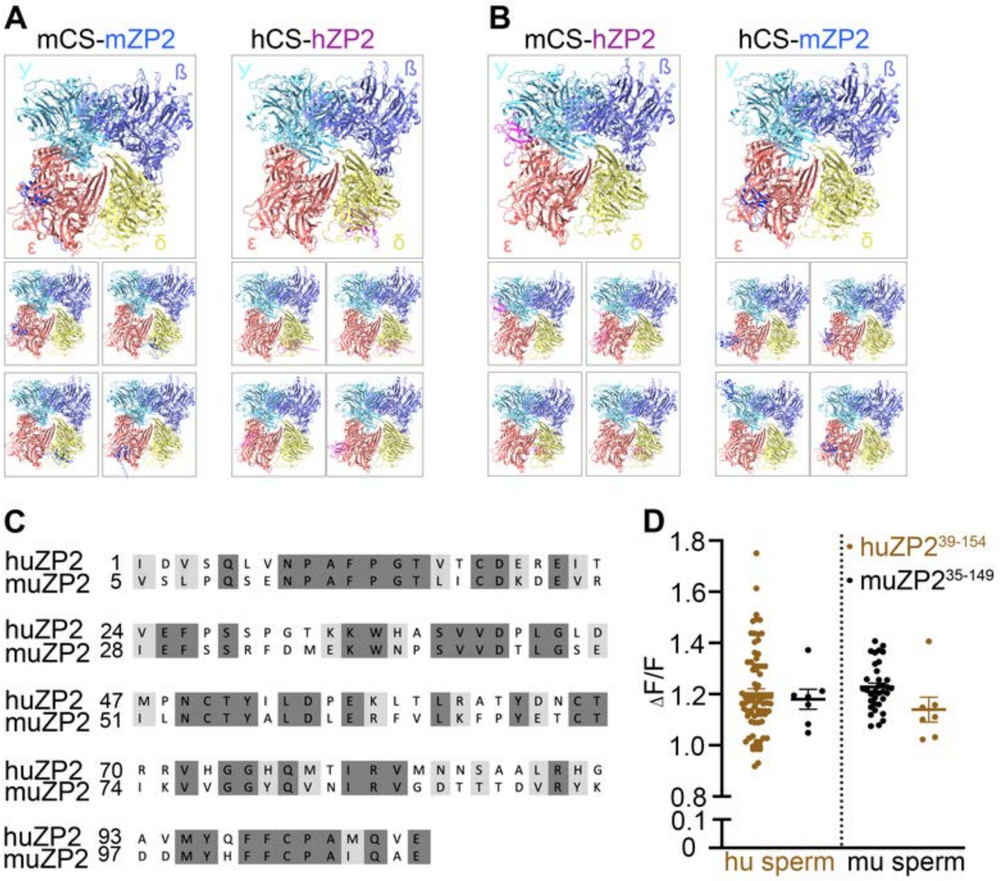


**Supplementary Fig. S4: Sequence similarity between the N-terminal muZP2$^{35\text{-}149}$ and huZP$^{239\text{-}154}$**

(A-B) Prediction models of the CatSper canopy and ZP2 from AlphaFold3 (Five models were generated from each set of input sequences). Intraspecies multimer prediction (A) and interspecies multimer prediction (B). muZP2$^{35\text{-}149}$ and huZP2$^{38\text{-}154}$ are colored in blue and magenta, respectively. Extracellular domains of CatSper are colored (β= light purple; γ= sky blue; δ= salmon; ε= yellow). (C) Amino acid sequence alignment of N-terminal huZP2$^{39\text{-}154}$ and muZP2$^{35\text{-}149}$. Identic amino acids (aa) were color-coded in dark grey and aa with similar chemical in bright grey. Analysis reveals 66% similarity (identical + similar aa). (D) Statistical analysis of the $[Ca^{2+}]i$ increase after stimulation with either huZP2$^{39\text{-}154}$ or muZP2$^{35\text{-}149}$ in murine or human spermatozoa. $N_{experiments} \geq 7$ experiments each with at least 6.25 Mio cells per ml.
